# Striking Long Term Beneficial Effects of Single Dose Psilocybin and Psychedelic Mushroom Extract in the SAPAP3 Rodent Model of OCD-Like Excessive Self-Grooming

**DOI:** 10.1101/2024.06.25.600634

**Authors:** Michal Brownstien, Michal Lazar, Alexander Botvinnik, Chloe Shevakh, Karin Blakolmer, Leonard Lerer, Tzuri Lifschytz, Bernard Lerer

## Abstract

Obsessive compulsive disorder (OCD) is a highly prevalent disorder that causes serious disability. Available treatments leave 40% or more of people with OCD significantly symptomatic. There is an urgent need for novel therapeutic approaches. Mice that carry a homozygous deletion of the SAPAP3 gene (SAPAP3 KO) manifest a phenotype of excessive self-grooming, tic-like head-body twitches and anxiety. These behaviors closely resemble pathological self-grooming behaviors observed in humans in conditions that overlap with OCD. Following a preliminary report that the tryptaminergic psychedelic, psilocybin, may reduce symptoms in patients with OCD, we undertook a randomized controlled trial of psilocybin in 50 SAPAP3 KO mice (28 male, 22 female). Mice that fulfilled inclusion criteria were randomly assigned to a single intraperitoneal injection of psilocybin (4.4 mg/kg), psychedelic mushroom extract (encompassing the same psilocybin dose) or vehicle control and were evaluated after 2, 4 and 21 days by a rater blind to treatment allocation for grooming characteristics, head-body twitches, anxiety and other behavioral features. Mice treated with vehicle (n=18) manifested a 118.71±95.96 % increase in total self-grooming (the primary outcome measure) over the 21-day observation period. In contrast, total self-grooming decreased by 14.60%±17.90% in mice treated with psilocybin (n=16) and by 19.20±20.05% in mice treated with psychedelic mushroom extract (n=16) (p=.001 for effect of time; p=.0001 for time X treatment interaction). 5 mice were dropped from the vehicle group because they developed skin lesions; 4 from the psilocybin group and none from the psychedelic mushroom extract group. Secondary outcome measures such as head-body twitches and anxiety all showed a significant improvement over 21 days. Notably, in mice that responded to psilocybin (n=12) and psychedelic mushroom extract (n=13), the beneficial effect of a single treatment persisted up to 7 weeks. Mice initially treated with vehicle and non-responsive, showed a clear and lasting therapeutic response when treated with a single dose of psilocybin or psychedelic mushroom extract and followed for a further 3 weeks. While equivalent to psilocybin in overall effect on self-grooming, psychedelic mushroom extract showed superior effects in alleviating head-body twitches and anxiety. These findings strongly justify clinical trials of psilocybin in the treatment of OCD and further studies aimed at elucidating mechanisms that underlie the long-term effects to alleviate excessive self-grooming observed in this study.

**Graphical Abstract:** 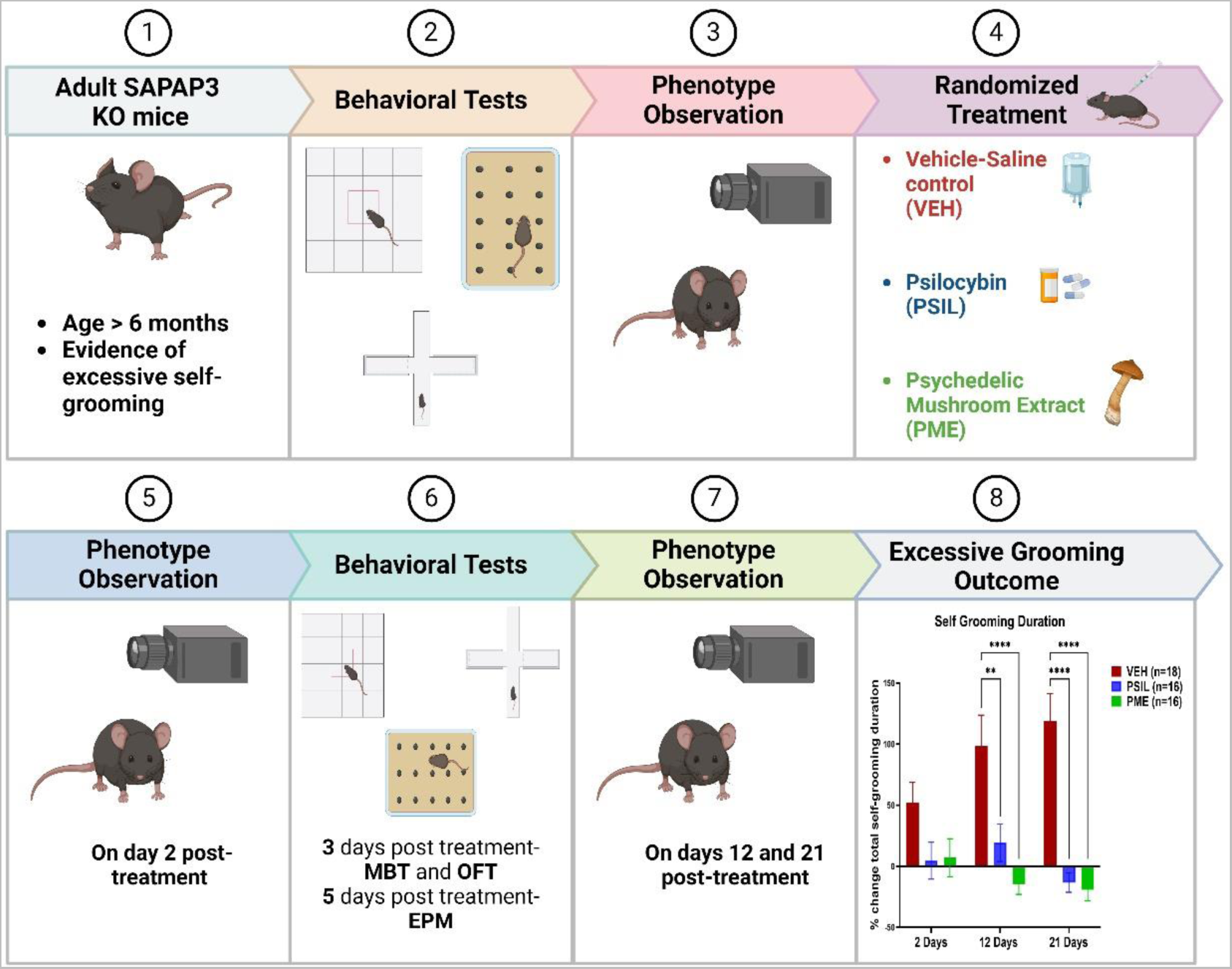

Prepared with BioRender (https://www.biorender.com/)

## 1. Introduction

Obsessive-Compulsive Disorder (OCD) affects 2-3% of the US population and is characterized by intrusive thoughts (obsessions) and repetitive actions (compulsions) ^1, 2^. The level of functional disability induced by OCD may be substantial. Exposure and response prevention (ERP) therapy is the most established cognitive-behavioral strategy for the treatment of OCD. Specific serotonin reuptake inhibitors (SSRIs) are the most widely used pharmacological treatment ^2^. However, 40–60% of patients experience only partial recovery and about 10% remain treatment resistant ^3^.

Pathological repetitive behaviors that characterize OCD also occur in disorders such Tourette Syndrome (TS), a childhood-onset neurodevelopmental disorder that manifests as tics and may be comorbid with OCD ^4^. Another disorder that overlaps with OCD is trichotillomania which is characterized by abnormal repetitive behaviors such as hair-pulling or skin-picking. Self-grooming, a normal phenomenon in rodents, occurs to a strikingly excessive extent in mice that carry a deletion of the SAP90/PSD95-associated protein 3 (SAPAP3) ^5^.

Synapse associated proteins (SAPAPs) are a family of proteins that are expressed in the brain and are highly concentrated in the postsynaptic density (PSD), an extensive complex associated with postsynaptic membranes of excitatory synapses ^6^.The SAPAP protein family can interact with postsynaptic molecules, including two multidomain scaffolding proteins - the SHANK protein family and PSD95, creating a regulator scaffolding complex at the excitatory synapse ^7^. This scaffolding complex anchors neurotransmitter receptors and signaling molecules to the postsynaptic membrane of the excitatory synapse and is responsible for the influence of SAPAP family on activity of ionotropic and/or metabotropic glutamate receptor ^8^. SAPAP3 (SAP90/PSD95 associated protein 3) is part of the scaffolding complex and is widely expressed in the striatum ^5^. SAPAP3 proteins have an important role in the function of postsynaptic glutamatergic synapses in the area ^5, 9^. Loss of these proteins leads to reduction of AMPA receptor-mediated corticostriatal excitatory transmission in medium-spine neurons (MSN), specific for the cortico-striatal circuit ^10^.

SAPAP3 KO mice begin to manifest the phenotype of excessive self-grooming at 4-6 months and the behavior increases in intensity in the following months frequently leading to severe skin lesions that are life-threatening. It has also been reported that SAPAP3 KO mice manifest increased anxiety ^5^, impaired cognitive flexibility ^11^ and aberrant habit formation ^12^. Fluoxetine treatment (5mg/kg, i.p. for 6 days) significantly reduced excessive grooming in SAPAP3 knockout (KO) mice and also anxiety-like behaviors without affecting total activity. A single injection of fluoxetine (5mg/kg, i.p.) did not affect the excessive grooming behavior of SAPAP3 KO mice. However, daily injections over 7 days led to a significant improvement in self-grooming behavior and anxiety ^5^. Ketamine has been reported to rapidly alleviate symptoms in patients with OCD ^13^. Recently, ketamine was found to attenuate self-grooming duration in SAPAP3 KO mice. A single ketamine injection significantly reduced grooming frequency at 1 hour and 1 day post-treatment but attenuated self-grooming was no longer observed by day 3 ^14^.

There is increasing interest in the use of psilocybin and other tryptaminergic psychedelics to treat a range of psychedelic disorders including major depression, alcohol and nicotine addiction and end of life anxiety^15^. In an open study, 9 patients with OCD were administered 29 doses of psilocybin ranging from sub-hallucinogenic to frankly hallucinogenic. Reductions in OCD symptoms were observed in all subjects and these generally lasted for at least 24 hours ^16^. Anti-OCD effects of psilocybin and psilocybin mushroom extract have also been reported in single patients (^17, 18^, in a pilot study of patients with serotonin reuptake inhibitor-resistant body dysmorphic disorder ^19^ and in mouse studies of marble-burying, a rodent behavior regarded as predictive of ant-obsessive effects ^20, 21^. The effect of psilocybin on excessive self-grooming and other phenotypes manifested by SAPAP3 KO mice is not known. A further question is whether entourage molecules contained in psilocybin mushroom extract might enhance a therapeutic effect of psilocybin in SAPAP3 KO mice ^22^.

In this study, we examined the effect of a single injection of synthetic psilocybin (PSIL) or psilocybin-containing psychedelic mushroom extract (PME) on excessive self-grooming, head-body twitches, anxiety and other behavioral phenotypes in SAPAP 3 KO mice, employing a randomized, controlled design with extended follow-up. We report that single administration of both treatments significantly attenuated the substantial increase in self-grooming observed in vehicle treated mice with surprising long-term effects.

## 2. Methods

### a. Animals

The SAPAP3 breeding colony was established using 5 heterozygous SAPAP3 KO male and female mice kindly provided by Dr Guoping Feng (Massachusetts Institute of Technology). We paired heterozygous male with heterozygous females at age 7-8 weeks, yielding offspring in the proportions of 25% wild-type, 25% homozygous and 50% heterozygous. Each mouse that is born in the reproduction colony goes through genotyping by the age of 4 weeks and is separated from the cage in which it was born. The mice are divided into separate cages by gender and age. Part of the mice are used for reproduction maintenance, part as a control group (wild type mice) and others used to study the SAPAP3 phenotype and the effects of possible treatments. Throughout the experiment male and female SAPAP3 KO mice were assessed separately in order to determine if there is significant difference in the behavioral phenotype between the genders.

Mice were housed under standardized conditions with a 12-hour light/dark cycle, stable temperature (22±1°C), controlled humidity (55±10%), free access to mouse colony chow and water and up to 8 per cage. Both female and male experimenters handled the mice and performed the studies. The study cages included both genotypes. Mice from the same cage received the same treatment.

Experiments were approved by the Authority for Biological and Biomedical Models, Hebrew University of Jerusalem, Israel (Animal Care and Use Committee Approval Number: MD-21-16596-4). All efforts were made to minimize animal suffering and the number of animals used. Each mouse was inspected before inclusion in the protocol for any signs of physical damage to the body from the excessive self-grooming phenotype (or rarely from other mice) and subsequently at each evaluation. Mice showing skin lesions were not included in experiments and those that developed skin lesions during the treatment period were removed from the study.

### b. Genotyping

Following the “HOTSHOT” method, genotype was determined by polymerase chain reaction

(PCR) of mouse tail DNA or by mouse ear hole DNA.

Three different SAPAP3 KO primers are used to distinguish the genotypes of wild type, heterozygous and Homozygous mice as follow:

**Primer F1** (5’ ATTGGTAGGCAATACCAACAGG 3’) and **Primer R1** (5’GCAAAGGCTCTTCATATTGTTGG 3’) identify wild-type allele (around 147 base pairs), while **Primer F1 and TK F2** (5’ CTTTGTGGTTCTAAGTACTGTGG 3’) identify KO allele (around 222 base pairs) (Welch et al, 2007))

### c. Treatments

PSIL was provided by USONA and was administered at a dose of 4.4 mg per kg. PME was provided by Back of the Yards Algae Sciences (BYAS). The dose of PME was adjusted so that each mouse received 4.4 mg per kg of psilocybin. Details of the chemical composition and purity of PSIL and PME used in this study have been published previously ^22^ and are provided here in the Supplementary Information. PSIL and PME, were dissolved in 100% saline (0.9 NaCl) solution, and a new solution was made on each injection day. For control treatment, saline vehicle (VEH) was administered. All treatments were administered by intraperitoneal (i.p.) injection. All groups were administered the same volume of injection 10 µl/g per mouse.

### d. Study Design

This was a three-arm, randomized, controlled trial that was conducted at the Hadassah BrainLabs Center for Psychedelic Research, Hebrew University, Jerusalem, Israel (Fig 1 – Consort Chart). Adult (6.5-7-month-old) SAPAP3 male and female mice that fulfilled inclusion criteria were randomly assigned by lottery to treatment with a single i.p. injection of PSIL, PME or saline vehicle (VEH). Random assignment was separate for male and female mice and treatment group size was constrained to insure a sex balance. The study began on 16.04.2023 and ended on 03.01.2024. Prior to the trial, adult (6-7 month) WT mice were compared to adult SAPAP3 KO mice to establish the existence of an excessive self-grooming phenotype. There were no significant differences in baseline characteristics among the groups (Table 1). Evaluations of grooming behavior and head body twitches were performed 48 hours, 12 days and 21 days after treatment by an observer (MB) who was blind to treatment assignment.

**Fig. 1:**
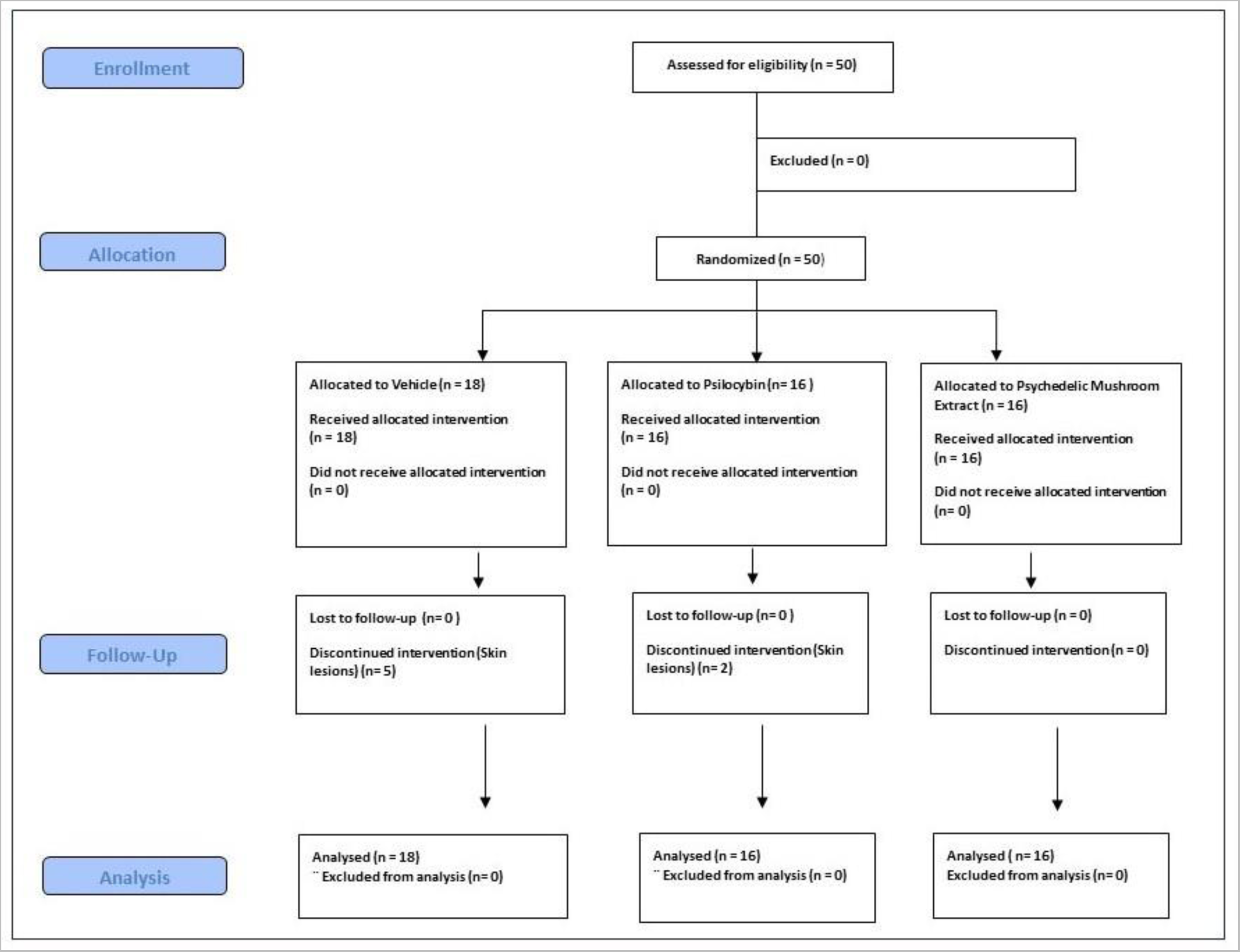
Consort chart showing the structure of the clinical trial.

**Fig 2:**
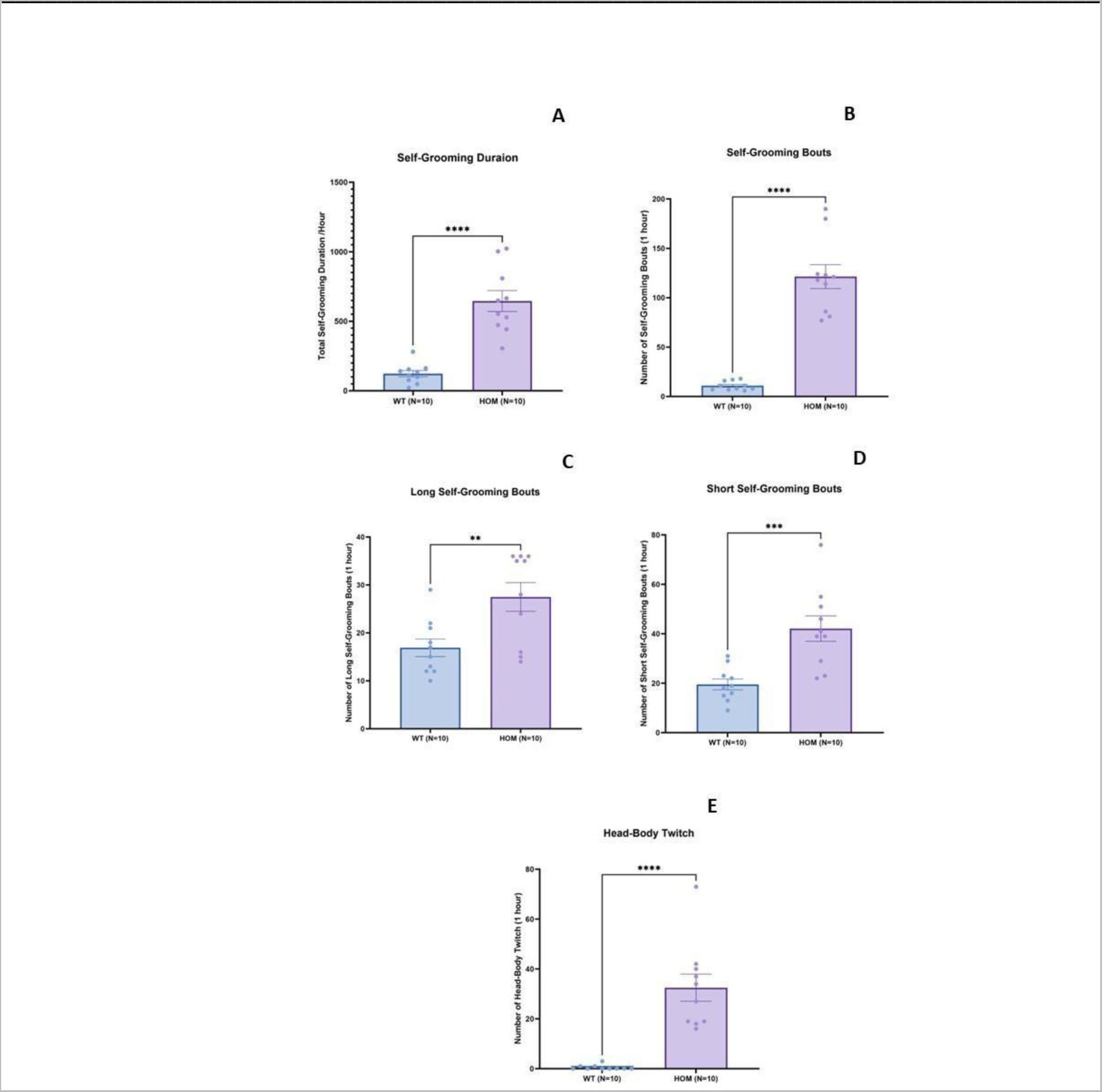
Comparison of self-grooming parameters (A-D) and head-body twitches [E] in SAPAP3 KO mice (n=10) and wild type littermates (n=10). Bars represent mean±SEM. **p<.01; ***p<.001; ****p<.0001 (Students t test, two tailed; unequal variance)

**Table 1:**
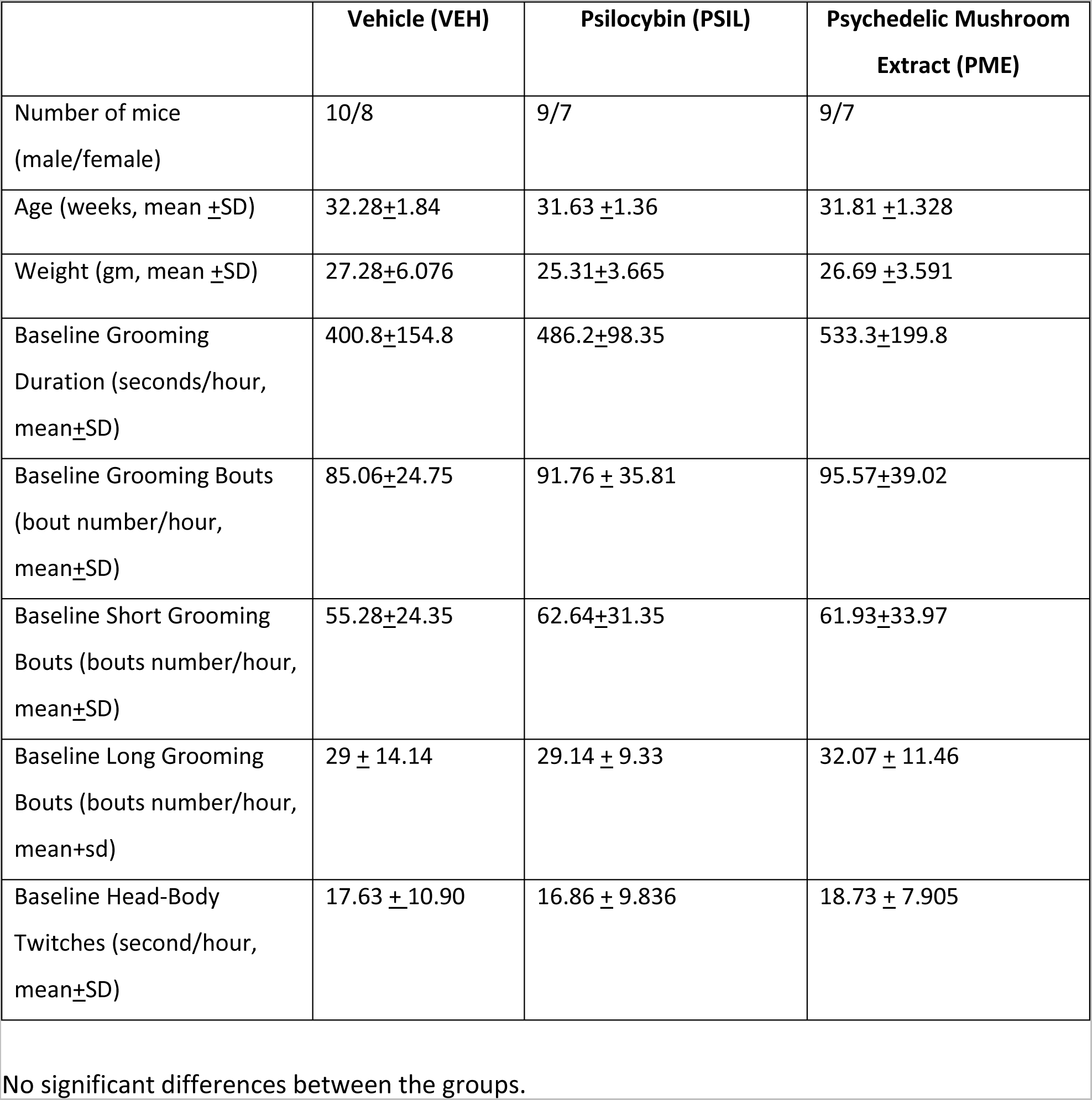
Background characteristics of the 3 treatment groups.

The pre-defined primary outcome measure of the study was percent change in total grooming score over one hour from the pretreatment score to 21 days as determined by analysis of variance with repeated measures. Secondary outcome measures were changes over the same treatment period in other grooming measures, head body twitches, anxiety levels and marble burying. Mice that dropped out of the study because of skin lesions were included in the analysis with their last observation carried forward.

After the 21-day assessment, a follow-up study was initiated in which mice were divided into the following groups according to their treatment in the study and the degree to which their self-grooming had improved. 1) Mice whose total self-grooming behavior over one hour had decreased by 10% or more from baseline (N=22) did not receive any further treatment and were reassessed 28 and 42 days following their initial injection. The distribution of these mice among the treatment groups was: VEH, N=0; PSIL, N=9; PME, N=13. 2). Mice in the VEH group whose total self-grooming score had worsened by more than 10% but had no skin lesions (N=15), were randomly assigned to treatment with PSIL (N=8) or PME (N=7) at the same dose as the mice in the PSIL and PME treatment groups received in the first injection. These mice then continued follow up of their self-grooming behavior which was evaluated 7 and 21 days after the second injection (28 and 42 days after the first [VEH] injection). The evaluator (MB) was not blind to treatment for assessments performed in the follow-up study. Mice that had been treated with PSIL or PME (N=6) at baseline and whose total self-grooming scores had improved by less than 10% from baseline or had worsened, were not followed up further.

### e. Inclusion and Exclusion Criteria

i. *Inclusion Criteria*
  1. Age 30-33 weeks
  2. Weight >18 gm.
  3. Genotype – SAPAP3 knockout
  4. Gender-male and females
  5. Total grooming time more than 300 seconds per hour
ii. *Exclusion Criteria*
  1. Presence of any skin lesion
  2. Loss of 10% body weight in two repeated measurements in the 4 weeks before the study
  3. Signs that the mouse had been the object of aggression

### f. Evaluation of Grooming Behavior

#### (i) Video Acquisition

Mice were temporarily separated from their littermates for a one-hour video recording session by a digital Go-Pro9 camera in a transparent plastic box (40 cm(l) x 60 cm ()( x 75 cm (h)), filled with regular wood bedding and the digital Go-Pro9 camera in the front. There was a half-hour of habituation to the setup prior to the video recording. The video recording was conducted in a SPF room, with lighting and average temperature of 22±1c between the hours of 8am-6pm. Between recordings the box was cleaned with ethanol and the bedding was changed.

#### (ii) Self-Grooming Behavior

Self-grooming is a natural and frequent phenomenon in rodents and other animals which includes a sequence of movements that aims to clean and maintain the fur. Self-grooming behavior includes face-wiping, licking, scratching and/or rubbing the head, snout and ears, and also includes full body grooming ^23^. The behavior of each mouse was characterized as follows: *Total Grooming Duration* - Total time of self-grooming in the course of one hour.

*Total Grooming Bouts* - The total number of self-grooming bouts in one hour.

A grooming bout was counted separately when there was more than two seconds difference between two bouts ^23^.

*Number of Short Grooming Bouts* **-** The total number of short self-grooming bouts under 3 seconds in one hour. A self-grooming was considered short if it lasted up to 3 seconds ^24^.

*Number Long Grooming Bouts* - The total number of long self-grooming bouts in one hour. A self-grooming bout was considered long if it lasted more than 3 seconds ^24^.

*Total Head-Body Twitches* - The total number of times, in one hour a mouse had a head-body twitch. This sudden, very rapid, repetitive, uncontrolled behavior is described as an axial move^25^ and is similar to human tics ^26^.

#### (iii) Video Analysis

All videos were scored by one rater (MB) who was blind to treatment assignment. In addition, a subset of 45 videos was scored by one of two additional raters. Cohen’s Kappas for comparison of total grooming scores between the additional raters and MB was 0.81.

#### (iv) Grooming Behavior in WT mice

10 adult WT mice, age 7 months, 5 females and 5 males, were used for the basic behavioral characterization. Each mouse was individually recorded for a one-hour video as described above and its self-grooming characteristics were analyzed according to the five categories described.

#### g. General Follow-up

i. *Weight: Each* mouse was weighed before entering the trial and twice a week thereafter.
ii. *Skin Lesions:* during the treatment protocol each mouse was tested visually to see if it developed skin lesions (on the head, neck, snout or body) due to excessive self-grooming. Mice that developed skin lesions were removed from the study.

#### h. Behavioral Tests

SAPAP3 KO (N=20) and WT (N=20) mice underwent a series of behavioral tests before entering the trial. These tests were performed 2--6 days before the initiation of the trial and then 3-5 days after treatment and included the following:

*Marble Burying Test (MBT)* was conducted to assess compulsive-like behaviors in HOM SAPAP3 mice compared to WT littermates, and to evaluate the effects of treatments on this behavior. The test was conducted in a quiet room with dim light, to minimize anxiety’s effect on behavior. A mouse was placed in a separate transparent cage with a cover, filled with 4.5 cm of new bedding shred and 20 glass marbles were evenly distributed in a 5x4 pattern (while keeping 2 cm distance between marbles and cage). After a 30 minutes test the number of covered marbles was determined by considering a marble buried if at least two-thirds of its surface was covered with bedding material ^27^.

*Open Field (OFT)* was performed immediately after the MBT to evaluate anxiety levels and locomotor activity and anxiety and the effects of treatments on this behavior. A mouse that spends more time in the periphery area is considered more anxious. The arena consisted of a square wooden arena (50 × 50 × 40 cm) with white walls and floor. In the beginning of the test a mouse was placed in the arena center, and was free to explore it for 10 minutes. Total duration of time (sec) spent in center arena, total duration of time (sec) spent in periphery arena and total distance traveled (cm) were measured by the Ethovision XT-12 Video Tracking System (Noldus Information Technology BV). Before starting a new test, the arena was cleaned with a 70% alcohol solution ^28^.

*Elevated Plus Maze (EPM)* was performed to evaluate the initial anxiety and the effect of treatments on this behavior. The test apparatus comprises four arms: two enclosed arms with high side walls and two open arms, all converging on a central platform elevated 75 cm above the ground. The animal is placed on the central platform and can explore the arena freely for 6-minutes. Ethovision XT-12 Video Tracking System (Noldus Information Technology BV) tracks the animals and concludes the total time spent in each arm. Before starting a new EPM test the arena was cleaned with a 70% alcohol solution ^29^.

### i. Statistical Analysis

The experimental data are presented as the mean ± standard deviation of the mean (SDM) in the text and as mean ± standard error of the mean (SEM) in figures. Inter-group differences were determined using one- and two-way repeated measures analysis of variance (ANOVA), as appropriate, and two-tailed t-tests. Sidak multiple comparison tests were used to analyze post-hoc comparisons. p<0.05 (two tailed) was considered statistically significant. Samples with values exceeding two standard deviations from the mean were excluded from the analysis. Grooming phenotype data was collected from 4 scorers on a subset of assessments and Cohen’s Kappa coefficient was performed on this data with confidence intervals of 95%. A Cohen’s Kappa score between 0.61-0.8 is considered substantial agreement and score between 0.8-1 is considered almost perfect agreement ^30^.

All statistical analyses were conducted using GraphPad Prism version 10.2.23.

## Results

### a. Self-Grooming and Head-Body Twitches in SAPAP3 KO Mice Compared to Wild Type Littermates

We first sought to confirm the presence of the over-grooming phenotype in SAPA3 KO mice. Taking into account short as well as long grooming bouts, mice homozygous for SAPAP3 KO (n=10) groomed a total of 605.30±196.60 sec. (mean ± SDM) over the observation period of one hour compared to wild type littermates (n=10) which groomed 123.30±71.82 sec. (t=6.0, p<0001, Fig.1A). The overgrooming phenotype manifested strongly in the total number of grooming bouts over one hour (67.10±30.43. vs. 10.90±4.50, t=9.0, p<0.0001, Fig.1B) and the number of short (42.10±16.30. vs. 19.50±6.90, t=4.052, p<0.001, Fig.1C) and long grooming bout (26.50±9.02 vs. 16.90±5.82, t=3.01, p<0.001, Fig.1D). There was also a significant difference in the number of head-body twitches between the SAPAP3 KO and wild type mice (9.00±7.67 vs. 0.50±0.97, t=6.66, p<0.0001, Fig.1E). Wild type mice manifested virtually no head-body twitches. There was no significant difference between male and female SAPAP3 KO mice on any of the grooming parameters or on head-body twitches (Supplementary Table 1).

### b. Effect of PSIL and PME on Self-Grooming and Head-Body Twitches in SAPAP3 KO Mice

We compared percent change from baseline to 21 days in total grooming time over an hour among the 3 treatment groups. Interim assessments were performed after 2 and 12 days. VEH-treated SAPAP3 KO mice (n=18) manifested a 118.71±95.96 % increase in total self-grooming over the 21-day observation period. In contrast, total self-grooming decreased by 14.60%±17.90% in mice treated with PSIL (n=16) and by 19.20±20.05% in mice treated with PME (n=16) (Fig. 3). ANOVA with repeated measures revealed a significant effect of time (F=5.61; df 3,135; p=0.001) and a significant interaction between time and treatment (F=10.65; df 2,45; p<0.0001). Post hoc tests comparing mice treated with PSIL or PME to VEH and to each other at each time point revealed a significant reduction in total self-grooming duration at day 12 for both PSIL (p=0.003) and PME (p<0.0001) compared to VEH, in addition to a significant increase in total self-grooming duration in the VEH group at day 12 (p=0.002) and 21 (p<0.0001) compared to 2 days post treatment(Fig 3).

**Fig. 3:**
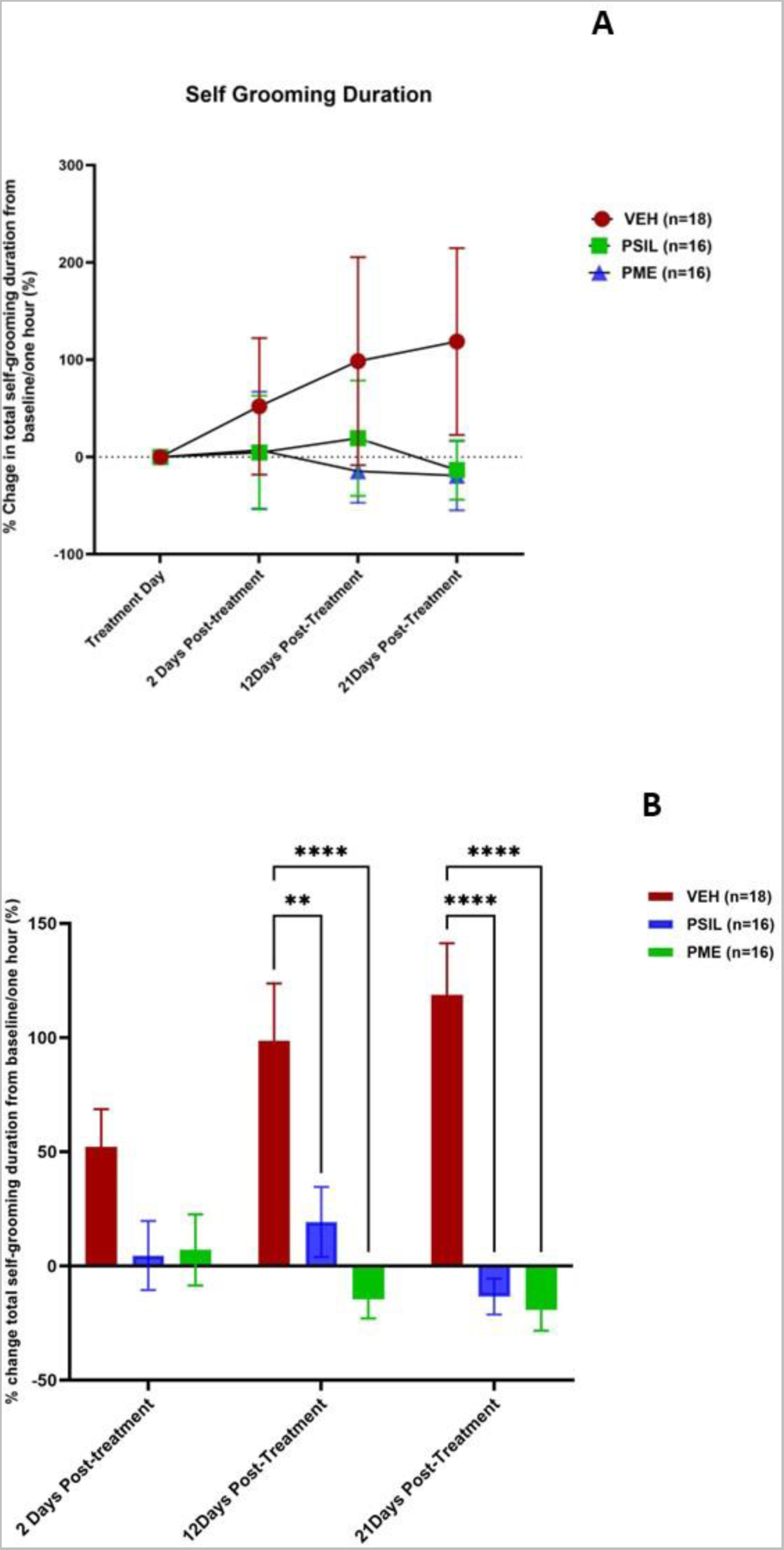
% Change in grooming duration over 21 days in SAPAP3 KO mice treated with VEH, PSIL or PME. Error bars represent SEM.. ANOVA for Panel A presented in the text. Panel B shows post-hoc comparisons (Sidak) of PSIL and PME to VEH. **p<.01; ****p<.0001 (versus VEH)

Similar results were observed for percent change from baseline in total number of grooming bouts over the course of 21 days of treatment. Vehicle treated SAPAP3 KO mice (n=18) manifested a 67.61±51.34% increase in number of self-grooming bouts over the 21-day observation period. In contrast, number of self-grooming bouts increased by only 4.28±27.14% in mice treated with PSIL (n=16) and decreased by 12.67±29.68% in mice treated with PME (n=16) (Fig. 4). ANOVA with repeated measures revealed a significant effect of time (F=3.854; df 3,141; p=0.01) a significant effect of treatment (F=6.317; df 2,47; p=0.003) and a significant interaction between time and treatment (F=5.92; df 6,141, p=<0.001). Post hoc tests revealed a significant reduction in self-grooming duration at day 12 for PME vs VEH (p=0.0152)(Fig 3). When long and short grooming bouts were analyzed separately, there was a significant effect of treatment with PSIL and PME to reduce long grooming bouts (F_treatmemt_=6.2; df 2,47; p=0.004). (Supplementary Fig. 2). Post-hoc tests (Supplementary Fig. 2) revealed a significant effect of PME after 12 days (p=0.04) and of both PSIL (p=0.001) and PME (p=0.001) after 21 days compared to VEH. In contrast, short grooming bouts increased under all treatments but not significantly. The increase with PSIL and PME was less striking than that observed in the VEH treated mice but differences were not statistically significant. Both PSIL and PME were associated with a significant reduction in head-body twitches over 21 days of treatment (F=3.939; df 2,47; p=0.026) (Fig 4). Post hoc testing revealed a reduction in head-body twitches at 12 days for both PSIL (p=0.01) and PME (p=0.01) (Fig. 4) versus VEH. There was no effect of treatments on the grooming phenotypes when comparing male and female mice (Supplementary Table 2)

**Fig. 4:**
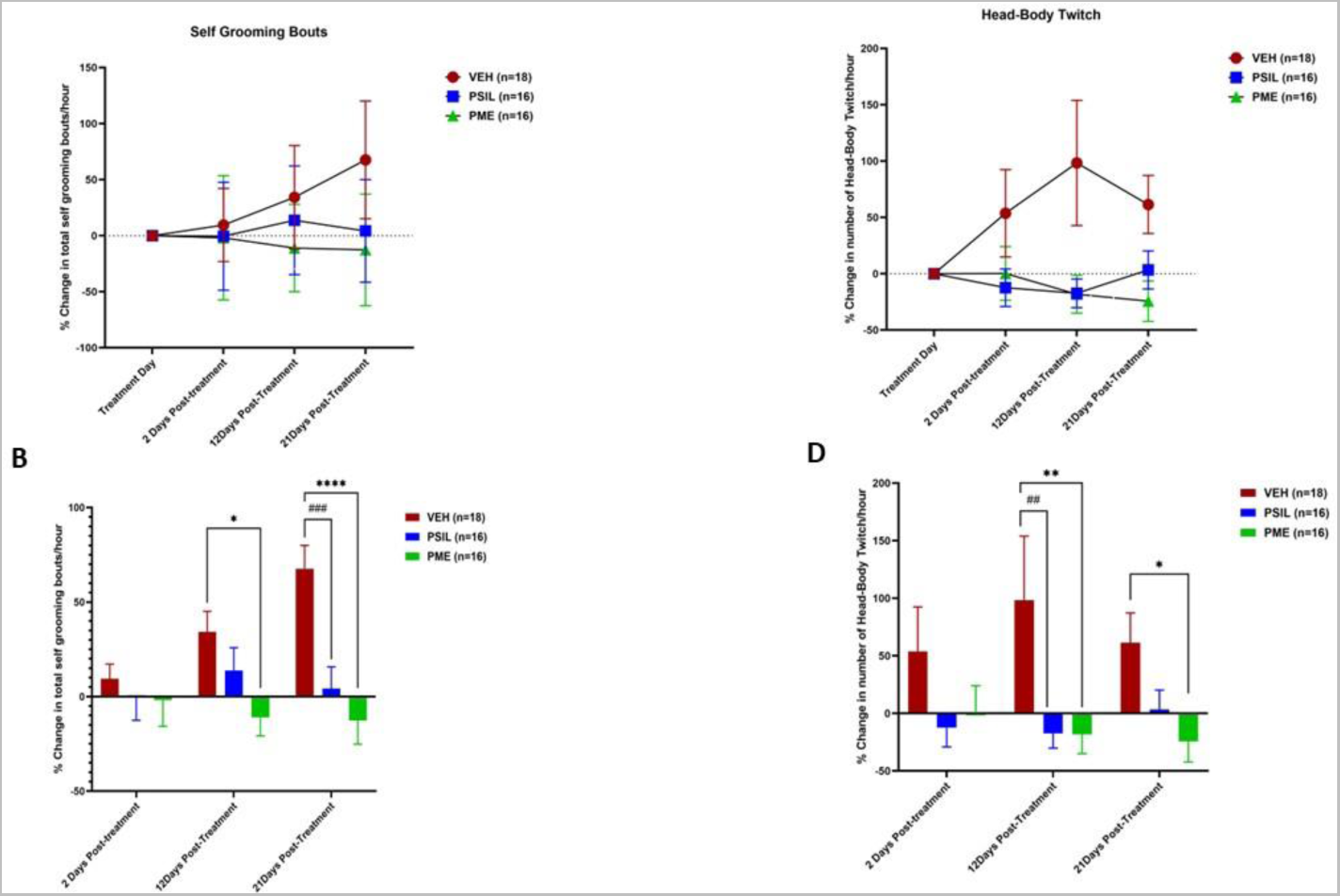
Self grooming bouts (A, B) and head body twitches (C,D) in SAPAP3 KO mice over 21 days following treatment with PSIL, PME or VEH. ANOVA with repeated measures for time course (A and C) presented in the text. Panel B shows post-hoc comparisons (Sidak) of PSIL and PME to VEH. *p<.05, **p<.01; ***p<.001; ****p<.0001 (versus VEH)

### b. Follow-up studies of the effect of PSIL and PME on excessive self-grooming

After 21 days, mice (PSIL n=12; PME n=13) whose self-grooming behavior had decreased by 10% or more from baseline were followed up without further treatment with further assessments of the self-grooming phenotype after 28 and 42 days. None of the mice who received VEH treatment met the improvement criterion and were thus not entered into the follow-up study. In evaluating the follow-up data for continued treatment after 21 days, it should be noted that these VEH-treated mice had an increase of 118,71±95.96% in their grooming scores after 21 days. 28 days after the treatment, in mice treated with PSIL percent change in total grooming duration over one hour was +8.25±49.17% compared to baseline and in mice treated with PME was −14.10±51.06%. 42 days after treatment in mice treated with PSIL total grooming duration over one hour was decreased by 5.2±56.32% from baseline and in mice treated with PME was reduced by 43.3±39.71%. Scores of total grooming duration over one hour show the consistent reduction in total grooming scores from baseline to 42 weeks in mice treated with PME as compared to the less consistent effect of PSIL with significant differences in favor of PME emerging at 12, 28 and 42 days post treatment (Fig. 5). Effects of PSIL and PME on head-body twitches show a similar pattern between the initiation of treatment and Day 42 (Fig 5).

**Fig. 5:**
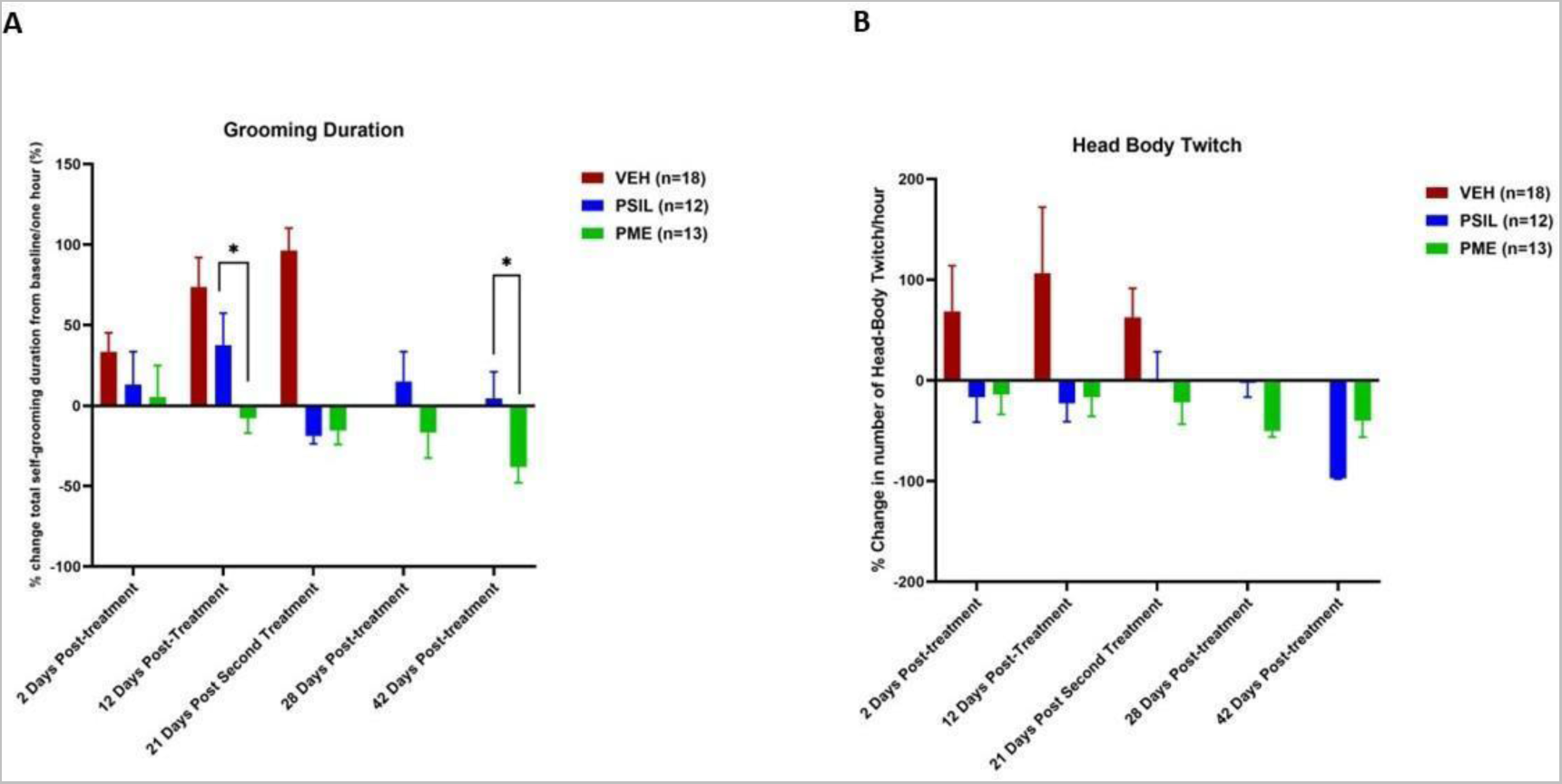
Treatment response from Day 21-42 of SAPAP3 KO mice that showed an improvement of 10% or more on total grooming score on Day 21. For total grooming score (A), ANOVA with repeated measures that included PSIL and PME groups only showed a significant effect of time (F=3.09; df 4, 76 ; p=.002). *p<.05, PME vs. PSIL on Days 12 and 42. (Sidak post hoc test). For head-body twitch, ANOVA with repeated measures that included PSIL and PME groups only showed a significant effect of time (F=4.61; df 4,80 ; p=.002) and time x treatment interaction (F=3.22; df 4,80 ; p=.01)

SAPAP3 KO mice treated with vehicle who reached Day 21 without developing skin lesions but did not show an improvement of 10% or more in total grooming scores from baseline, were randomised to receive a second injection of PSIL (12) or PME (13) at the same dose as the initial treatment. These mice were reevaluated for grooming parameters on days 28 and 42, 7 and 21 days respectively after receiving the second injection (Fig.6). Data were analyzed by ANOVA with repeated measures comparing percent change from day 21 when the second injection was given. Treatment response of mice treated with PSIL and PME was compared. There was a significant effect time (F=102.3; df 2.26; p<0.0001), total number of grooming bouts (F=55.46; df 2,26; p<0.0001) and number of head-body twitches (F=13.08; df 2,26; p=0.0001), with PSIL (p<.0001) and PME (p<.0001) both showing significant effects for the grooming parameters but only PSIL (p=0.046) for head-body twitches (Fig. 6).

**Fig. 6:**
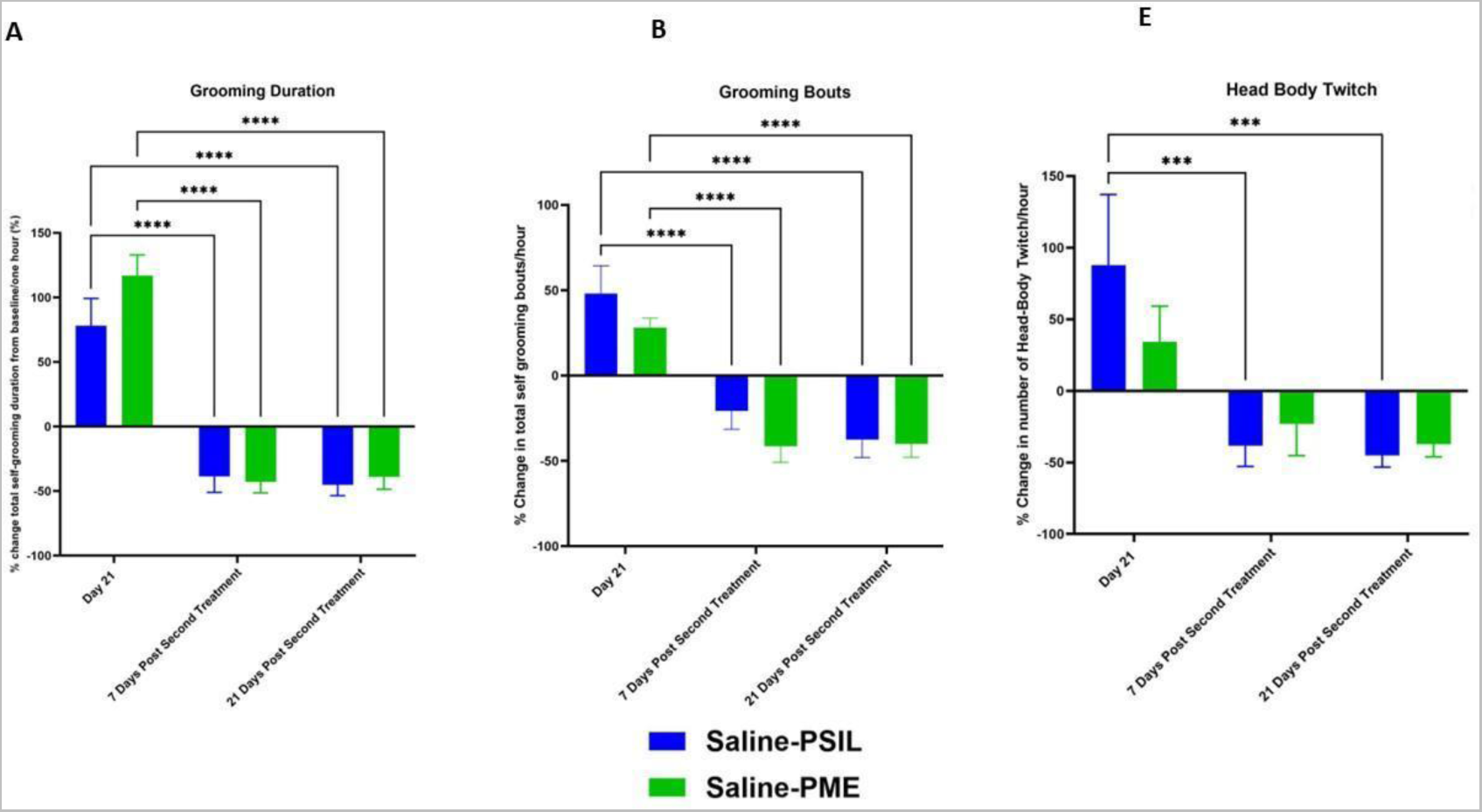
Effect of treatment with PSIL or PME on % change from Day 21 in grooming parameters (A and B) and head-body twitch [C) in SAPAP3 KO mice treated on Day 0 with VEH (saline) and on Day 21 with PSIL or PME. For total grooming duration (A) and grooming bouts (B) ****p<,0001 vs. Day 21. For head-body twitches (C) ***p<.001 vs. Day 21

### e. Behavioral Tests in SAPAP3 KO Mice Compared to Wild Type Littermates

We compared 20 SAPAP3 KO mice to 20 wild type littermates on a series of behavioral tests putatively reflecting activity, anxiety and compulsive behavior. On the open field test, there was no significant difference between the two groups in distance traveled (Fig7A). Time spent in the center of the open field was significantly lower in the SAPAP3 KO mice than in the wild type mice (Fig 7B), reflecting a higher level of anxiety in the KO mice. This higher anxiety level was also reflected in the elevated plus maze results in which KO mice spent significantly less time in the open arms (Fig. 7C) than their wild type littermates. Finally, on the marble burying test, SAPAP3 KO mice buried strikingly fewer marbles than their wild type littermates (Fig. 7D).

**Fig. 7:**
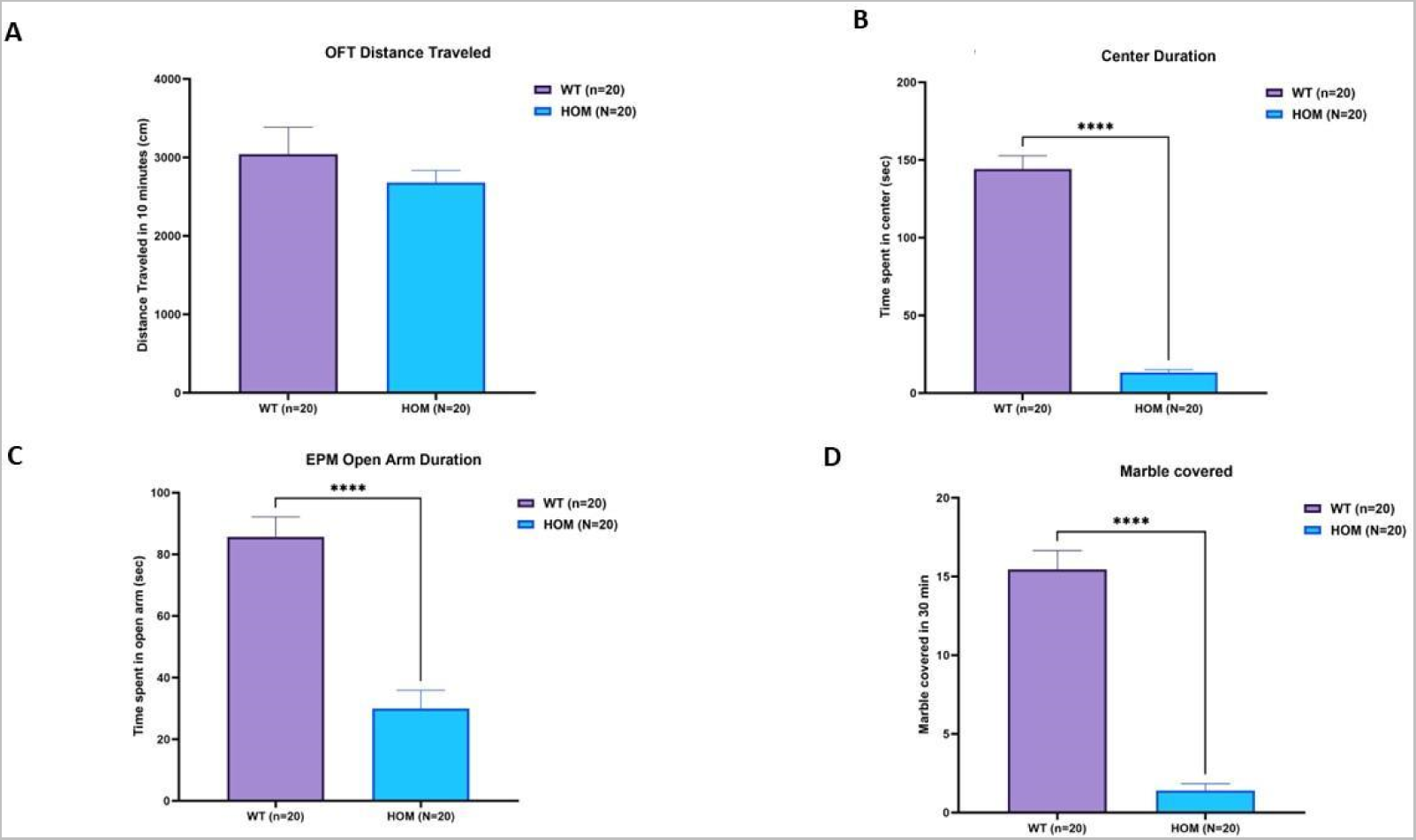
Open field distance travelled (A), Center Duration (B), EPM Open Arm duration (C) and Marble Burying in Wild Type (WT, n=20) versus SAPAP3 KO mice (HOM, n=20). ****p<.0001 (Students t test, two-tailed, unequal variance)

### f. Effect of PSIL and PME on Behavioral Tests in SAPAP3 K0 Mice

SAPAP3 KO mice underwent behavioral testing following treatment according to the following schedule: MBT and OFT 3 days; EPM 5 days following treatment. On the OFT, there was a significant interaction between time and treatment (F=5.22, df 2,47, p=0.009). On post hoc testing (Sidak) the time spent in the center of the open field was significantly greater following treatment in mice that received PME compared to mice that received VEH (p=0.019) or PSIL (p=0.009) (Fig 8A). As regards time spent in the periphery there was no difference between the groups before or after treatment. On the elevated plus maze there was a significant effect for time spent in the open arms before and after treatment (F=6.64; df 1,47; p=0.013). On post-hoc testing, time spent in the open arms was significantly increased for mice treated with PME (p=0.025). (Fig. 6C). There were no significant differences in time spent in the closed arms of the EPM. An intriguing finding that emerged from these analyses was for marble burying. Marble burying was strikingly increased by both treatments and two-way ANOVA revealed a highly significant effect of time (F=60.04, df 1,47 p<0.0001) and treatment (F=11.4, df 2,47 p<0.0001) and a significant time X treatment interaction (F=14.3, df 2,47 p<0.0001) (Fig. 6D). Post hoc testing showed a significant effect of PSIL (p<0.0001) and PME (p<0.0001) to increase marble burying and a significantly greater effect of PME compared to PSIL (p=0.038).

**Fig. 8:**
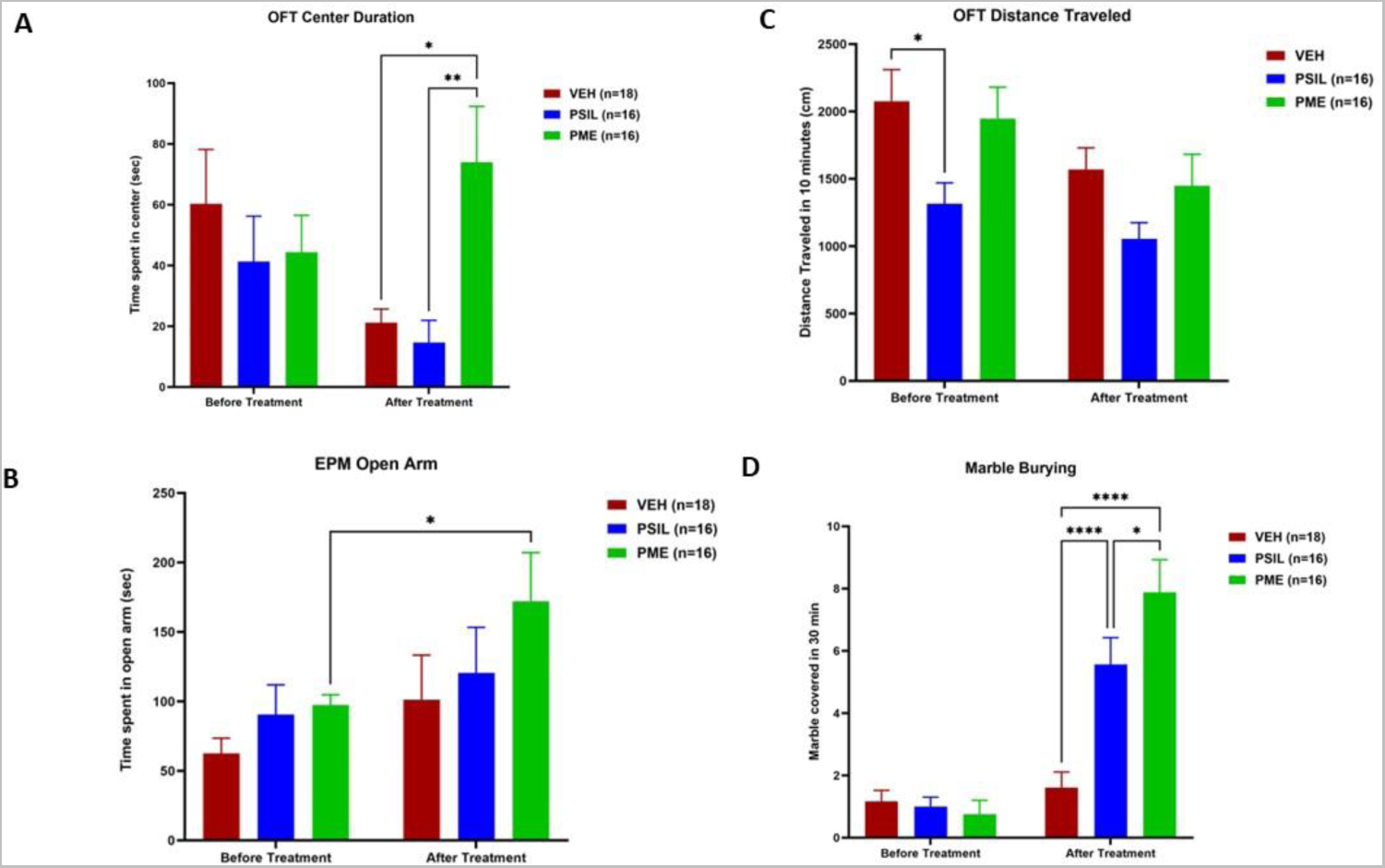
Open Field Test center duration (A) Open Field distance travelled (B) (C) Elevated Plus Maze open arm duration (C) and Marble burying (D), in SAPAP3 KO mice before and after treatment with VEH, PSIL or PME. ANOVA with repeated measures results given in the text. Post-hoc Sidak tests: *p<.05, **p<.02, ****p<.0001

### f. Adverse Events

7 mice developed skin lesions as a result of excessive self-grooming. They were withdrawn from the study but included in the analysis with their last observation carried forward. 5 were in the VEH group and 2 in the PSIL group. No mice in the PME group developed skin lesions. There were no instances of weight loss that required withdrawal from the study and no instances of attacks by mice on cage mates that resulted in obvious injury.

## Discussion

Examining male and female mice homozygous for deletion of the SAPAP3 gene, we have confirmed that these mice manifest excessive self-grooming and head-body twitches compared to wild type littermates as well higher levels of anxiety. Implementing a randomized, controlled trial in male and female SAPAP3 KO mice, we have shown that a single administration of synthetic psilocybin or psilocybin-containing psychedelic mushroom extract, attenuates the further increase in self-grooming and head-body shakes that is observed in vehicle treated mice and reduces anxiety. In mice that responded to the treatment, the beneficial effect on self-grooming and head-body twitches was maintained up to 6 weeks after a single injection of PSIL or PME. In an additional follow-up study, we demonstrated positive effects of PSIL and PME on the SAPAP3 KO phenotype in mice that had not responded to vehicle treatment in the main study with beneficial effects lasting for 3 weeks. The therapeutic effect of PSIL and PME was observed in male as well as female mice. These striking findings have significant translational implications for the treatment of OCD.

The compulsive component of OCD is characterized by repetitive behaviors over which the sufferer has limited control. The excessive self-grooming observed in SAPAP3 KO mice is a similar repetitive behavior that the mouse continues to perform in spite of deleterious consequences. SAPAP3 KO mice manifest a grooming disorder of extreme severity. SAPA3 KO mice also manifest significant anxiety which is frequently observed in patients with OCD. Grooming disorders such as trichotillomania, hair pulling and skin picking that occur in humans, are closely related to OCD and are classified under obsessive-compulsive and related disorders in the Diagnostic and Statistical Manual of Mental Disorders (DSM-5) ^31^. Tourette’s syndrome (TS) is a related disorder that is characterized by repeated, involuntary, motor and vocal tics ^4^. The head-body twitches that are observed in SAPAP3 KO mice are similar to the motor tics observed in patients with TS. Moreover, there is significant comorbidity between TS and OCD. Supporting a possible relationship between SAPAP3 and OCD in humans, there is evidence that polymorphisms in the SAPAP3 gene are associated with susceptibility to grooming disorders such as trichotillomania, pathological nail biting and skin picking in ^32^. Furthermore, in a sequencing study, rare variants in the human SAPAP3 gene were associated with trichotillomania and OCD ^33^. Thus, there is considerable support for the translational impact of our current findings which suggest that PSIL and PME may be effective treatments for OCD in humans and may be characterized by unique, long-acting effects of a single treatment.

In a recent study of ketamine in SAPAP3 KO mice, positive effects were observed in alleviating excessive grooming ^14^. However, these effects were no longer evident 3 days after the treatment whereas in the current study therapeutic effects of PSIL and PME persisted for up to 42 days following a single injection. This suggests that psilocybin, whether in chemical form or as a component of psychedelic mushroom extract, could be a unique, long-acting treatment for OCD. It is noteworthy that in depression as well, psilocybin has been reported to have considerably longer lasting positive effects than ketamine ^15^.

An interesting question raised by our findings Is whether entourage molecules found in psilocybin-containing psychedelic mushroom extract might enhance the therapeutic effect of psilocybin in SAPAP3 KO mice. In a recent study in which we measured synaptic protein levels after synthetic psilocybin and psilocybin-containing mushroom extract administration, we found that the extract had a more potent and prolonged effect on synaptic plasticity than synthetic psilocybin. In the same study, our ex vivo brain metabolomics data supported a gradient of effects from inert vehicle via chemical psilocybin to PME further supporting differential effects ^34^. In the current study the effect of PME was more pronounced than that of PSIL in certain cases, such as alleviating head-body twitches after 21 days and excessive grooming after 42 days. In relieving anxiety related phenotypes, the advantage of PME over PSIL was clearly evident in tests such as OFT (center duration), EPM (open arms) and marble burying.

The question of psilocybin dose merits consideration. In the current study we administered 4.4 mg/kg. Based on allometric calculations extensively discussed in our previous paper ^34^ this dose is approximately equivalent to 25 mg in a 70 kg person. It induces significant head twitch response in mice ^22^ and a strong psychedelic experience in humans ^15^. In the only clinical trial of psilocybin in OCD publishes to date, ^16^ observed that lower non-psychedelic doses than this as well as higher doses, induced improvement in OCD features. However, it should be noted that ^16^ reported only acute effects extending at most to 24 hours. Thus, the dose of psilocybin required in human studies to achieve long lasting anti-OCD effects remains an open question.

A further observation of interest in this study is that SAPAP3 KO mice manifested significantly lower levels of marble burying than wild type mice. In prior studies, psilocybin has been observed to reduce marble burying in mice ^20, 21, 27^, as do other treatments effective in treating OCD such as serotonin reuptake inhibitors. The lower level of marble burying was not a consequence of lower activity since there was no significant difference in distance moved on the open field test between SAPAP3 and wild type mice. There was a striking effect of treatment to enhance marble burying in the SAPAP3 KO mice. PSIL and PME both induced a significant increase in marble burying with the effect of PME being significantly stronger. Thus, we have observed that mice characterized by a phenotype that has high face validity for OCD and OCD-like disorders in humans and is alleviated by treatments that are effective in OCD, manifest marble burying behavior that is diametrically opposite to that observed in wild type mice. We speculate that the improved marble burying seen after treatment is a consequence of lower anxiety levels following treatment and a greater propensity to explore the cage, as seen is in the open field test, and in so doing engage in normal digging behaviors. Our finding need not detract from the validity of the MBT as a predictive test for anti-obsessive potential but it does further call into question the validity of this test as a model for OCD which has been questioned previously ^35^.

The mechanism whereby psilocybin attenuates excessive self-grooming and anxiety in mice and might attenuate OCD symptoms in humans is unclear at this time. Neuroimaging studies in OCD patients suggest that the disorder is associated with dysfunction of fronto-striatal circuits that play a role in cognitive flexibility and emotional regulation ^36^. ^14^ suggested that increased activity of dorsomedial prefrontal neurons projecting to the dorsomedial striatum may be implicated in the effect of ketamine to reduce excessive self-grooming in SAPAP3 KO mice. Studies aimed at determining whether an effect on fronto-striatal pathways plays a role in the effect of psilocybin on self-grooming in SAPAP3 KO mice is indicated. There is increasing evidence that glutamatergic mechanisms are implicated in OCD ^37^ and a role in excessive self-grooming by SAPAP3 KO mice is supported by short-term alleviation of this phenotype by ketamine ^14^ and by reports of ketamine efficacy in patients with OCD ^13^. Glutamatergic pathways may be may be implicated in the therapeutic effects of psilocybin in neuropsychiatric disorders ^38^ and may play a role in therapeutic effects in OCD and in alleviating the OCD-like phenotype of SAPAP3 KO mice. It has been shown that SAPAP3 deletion reduces AMPA-type glutamate receptor (AMPAR) - mediated synaptic transmission in striatal medium spiny neurons through postsynaptic endocytosis of AMPARs ^10^. A key question to be addressed is how psilocybin might impact on this mechanism.

In conclusion, we have shown that a single treatment with PSIL or PME significantly attenuates excessive self-grooming and reduces anxiety in SAPAP3 KO mice with the beneficial effect of treatment extending up to 6 weeks following a single administration. This finding provides strong translational support for trials of psilocybin and related tryptamine in OCD and related disorders in humans and a strong impetus for further preclinical and translational studies to understand the therapeutic mechanisms underlying these effects.

## Acknowledgments

Hodaya Denekamp and Noa Shatsnan assisted with ratings of videos.

## Author Contributions

Conception and design: BL, TL

Acquisition of data: MB, AB, CS

Analysis and interpretation of data: MB, AB, KB, LL, BL, TL

Drafting the article or revising it critically for important intellectual content: MB, AB, KB, LL, BL, TL

Final approval of the version to be published: All authors

## Data Availability

Data will be provided for further analysis to qualified investigators. Please contact

