## Supplemental Tables, Figures for "Striking Long Term Beneficial Effects of Single Dose Psilocybin and Psychedelic Mushroom Extract in the SAPAP3 Rodent Model of OCD-Like Excessive Self-Grooming"

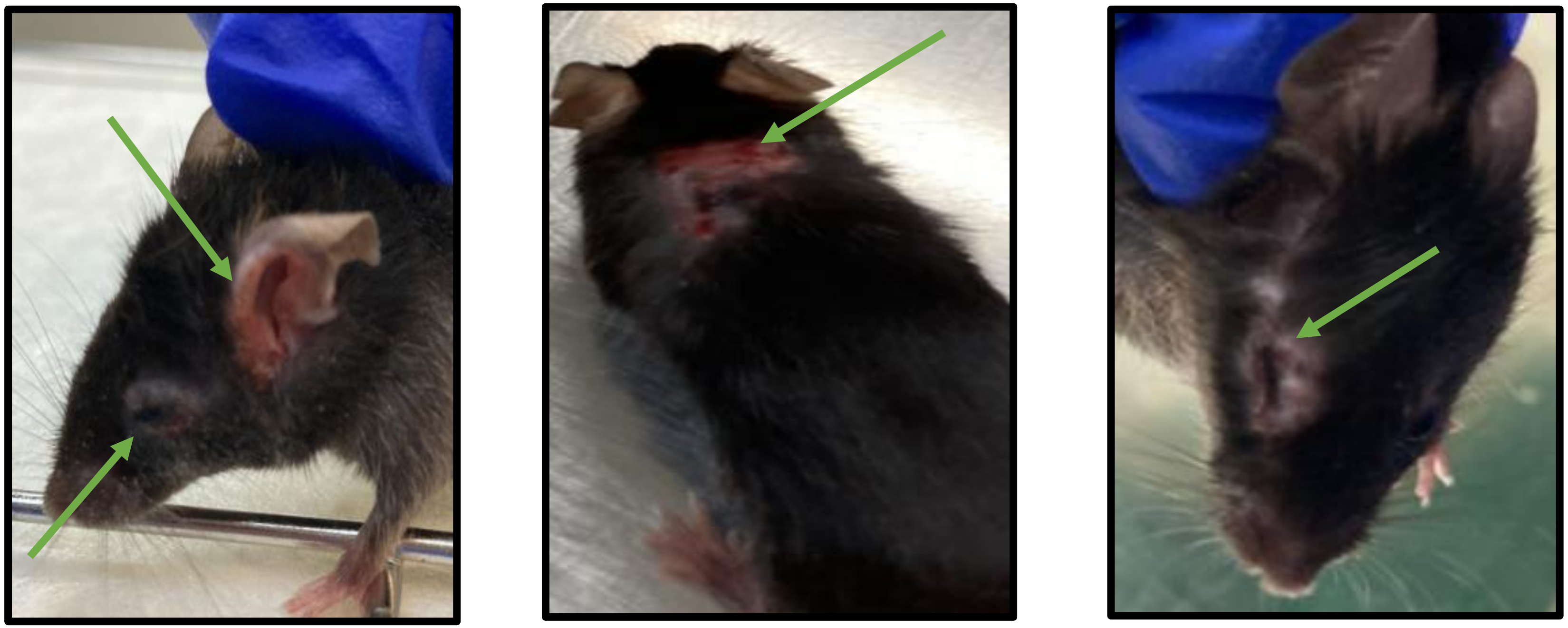

**Supplementary Figure 1:** SAPAP3 KO mice from the Vehicle-Saline group that developed skin lesions during the experiment. The lesions are around the eyes, ears and back area. These lesions are a result of excessive self-grooming.

**Supplementary Video 1**

Excessive Self Grooming in SAPAP3 KO Mice - Sapap3 KO Mice- Excessive self grooming phenotype (Click to view)

**Supplementary Table 1**: Baseline characteristics of female and male SAPAP3 KO mice and baseline grooming parameters

| | **Female** | **Male** |
|---|---|---|
| Number of mice | 5 | 5 |
| Age (weeks, mean ±SD) | 31.40 ±1.14 | 31.6 ±1.517 |
| Weight (gm, mean±SD)* | 20.40 ±2.302 | 29.20 ±2.864 |
| Baseline Grooming Duration (second/hour, mean+SD ) | 569.2 ±184.1 | 721.4 ± 279.6 |
| Baseline Grooming Bouts (bout number/hour, mean±SD ) | 104.2 ±23.15 | 138.6 ± 44.64 |
| Baseline Short Grooming Bouts (bout number/hour, mean±SD ) | 45. 8 ± 19.84 | 38.40 ± 12.84 |

| | | |
|---|---|---|
| Baseline Long Grooming Bouts (bout number/hour, mean±SD ) | 23.40 ± 8.649 | 31.60 ± 9.290 |
| Baseline Head-Body twitch (number/hour, mean±SD ) | 6.60 ± 4.519 | 5.40 ± 4.669 |

*There is a significant difference between the weight of female and males (p=0.0007)

**Supplementary Table 2:** Effect of treatment over 21 days on grooming parameters in male and female SAPAP3 KO mice

| | **Gender** | **Vehicle** | **Psilocybin** | **Psychedelic Mushroom Extract** |
|---|---|---|---|---|
| Grooming Duration (seconds/hour, mean+SD % change) | Male | 108.6 + 57.07 | -19.28 + 18.44 | -17.75 + 38.36 |
| | Female | 132.3 + 130.1 | 3.97 + 37.11 | -21.26 + 31.59 |
| % Change in Grooming Bouts (bout number/hour, mean+SD % change)) | Male | 26.20 + 22.99 | -13.87 + 21.37 | -10.26 + 51.93 |
| | Female | 58.02 + 48.38 | 27.54 + 545.66 | -15.77 + 18.75 |
| Short Grooming Bouts (bout number/hour, mean+SD % change)) | Male | 15.43 +25.30 | -17.17 + 29.47 | -30.59 + 43.04 |
| | Female | 68.53 + 58.11 | -13.04 + 50.61 | -41.59 + 19.02 |

| | | | | |
|---|---|---|---|---|
| Long Grooming Bouts (bout number/hour, mean±SD % change) | Male | 50.55 ± 48.88 | -19.78 ± 19.69 | -6.926 ± 55.15 |
| | Female | 106.3 ± 114.7 | 3.01 ± 62.74 | -12.91 ±27.96 |
| Head-Body twitch (second/hour, mean±SD % change)) | Male | 39.08± 63.28 | -12.44 ± 55.03 | -6.94 ± 90.25 |
| | Female | 83.39± 149 | 23.74 ± 80.73 | -46.80± 31.77 |

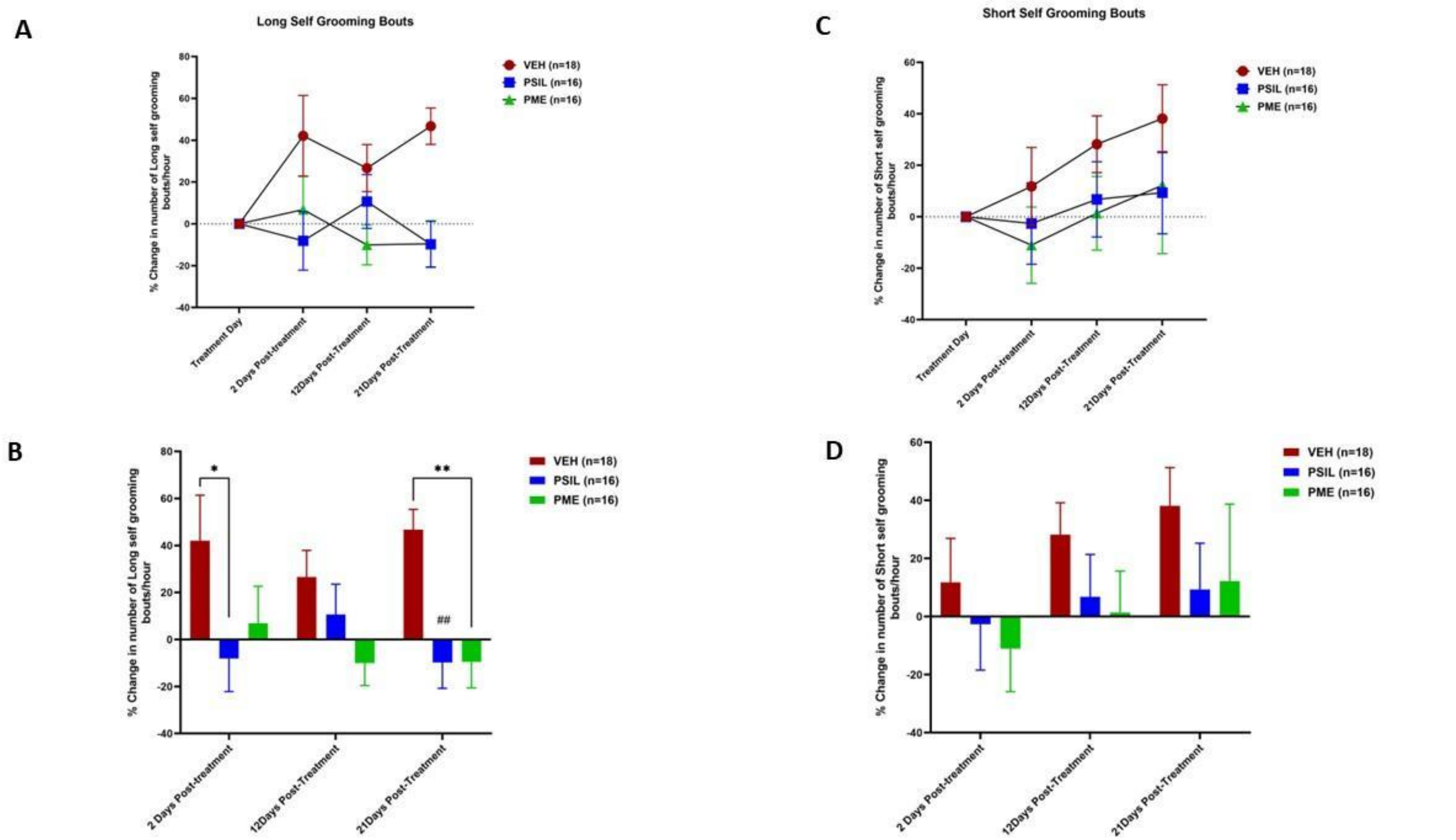


**Supplementary Fig. 1:** Effect of PSIL, PME and VEH on long and short grooming bouts in SAPAP3 KO mice. ANOVA with repeated measures on long grooming bouts showed a significant effect of treatment ($F=6.2$; df;2,47 $p=0.004$). *$p<.05$, **$p<.01$ (Sidak)

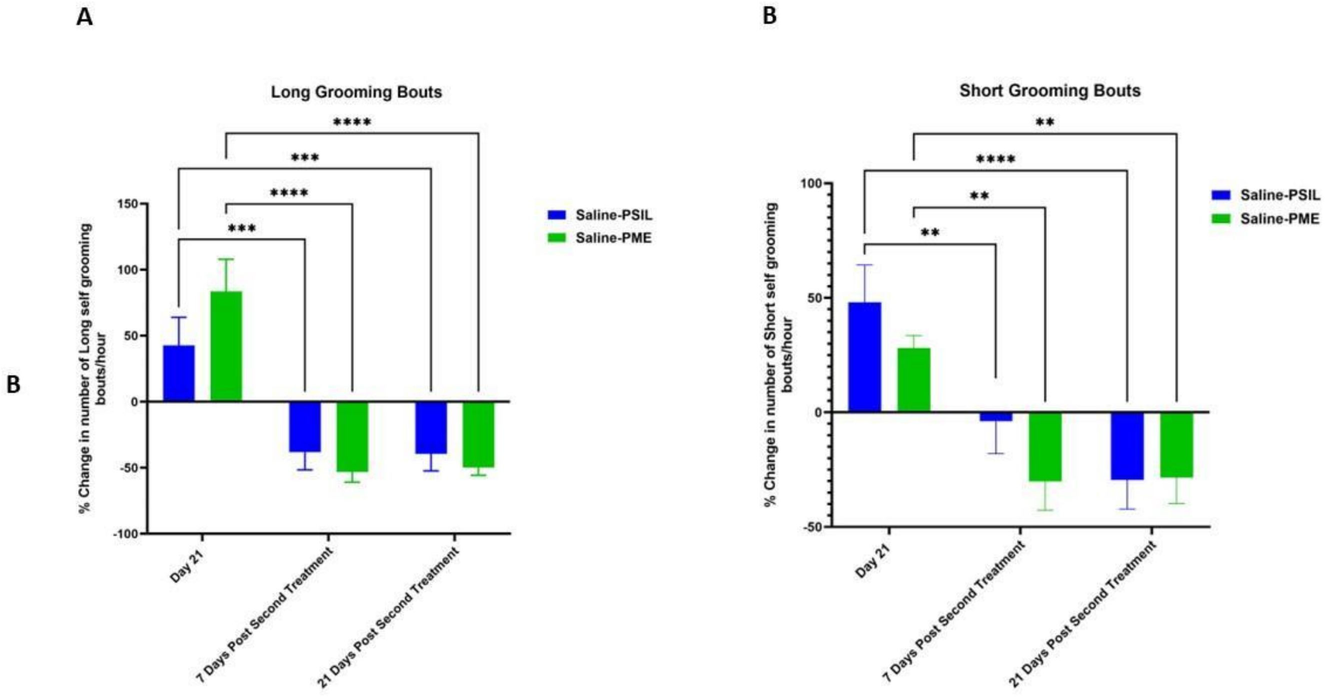


**Supplementary Fig. 2**: Effect on long (A) and short grooming bouts of treatment with PSIL or PME administered on Day 21 to SAPAP3 KO mice that had received VEH (saline) in the first part f the trial and did not show an improvement of 10% or more after 21 days. **p<.01, ***p<.001, *P<.0001 (Sidak's post hoc test)

**Chemical Composition and Purity of PSIL and PME**

PSIL was supplied by Usona Institute, (Madison WI, USA) and was determined by AUC at 269.00 nm (UPLC) to contain 98.75 wt. % psilocybin. PME was supplied by Back of the Yards Algae Sciences - Parow Entheobiosciences (Chicago, IL, USA) and was determined to contain psilocybin 1.3 wt.%, psilocin 0.17 wt.%, norpsilocin 0.01 wt.%, baeocystin 0.02 wt.%, norbaeocystin 0.007 wt.% and aerugeniscin 0.008 wt.% by LC-MS/MS (See section, PME cultivation, extraction and analysis, for description of analytic method). PSIL and PME were dissolved in 0.9% NaCl. Whether administered PSIL or PME, mice received psilocybin at a dose of 4.4 mg/kg intraperitoneally (i.p.). The dose of psilocybin that we used (4.4 mg/kg) was chosen on the basis of our previous dose response study on the effect of psilocybin on the head twitch response [28], and in view of the fact this dose is equivalent in mice to a 25 mg dose in humans according to the widely used DoseCal dose conversion methodology [34]. For PME the dose was based on the following calculation: To prepare a solution for a mouse that weighs 30 g so that the mouse will receive 4.4 mg/kg of pure psilocybin, $0.03\ kg\ (since\ 30\ g\ =\ 0.03\ kg)\ \times 4.4\ mg/kg = 0.132\ mg$ of psilocybin per 30 g mouse is needed. Since 1 mg of psilocybin is found in 76.92 mg of PME (1.3%) the amount of PSIL that was given needs to be multiplied by a factor of 76.92, $76.92 \times 0.132\ mg = 10.15\ mg$ of PME is needed per 30 g mouse. In all cases, injections were administered in a standard injection volume of 10 µl/g per mouse. Both PSIL and PME rapidly dissolved completely in saline (0.9% NaCl).

**PME cultivation, extraction and analysis**

*PME cultivation and extraction: Psilocybe cubensis* was cultivated from genetically confirmed spore material (see www.Entheome.org for whole-genome sequence) germinated on light malt extract agar in a petri dish. Agar-cultured colonies were transferred to sterilized grain jars (rye, white millet, and oats). The colonized grain was transferred into pasteurized bulk substrate bags. The mushrooms were fruited and harvested after 18-28 days. The biomass was dried at 65°C for 6 hours. The dried ground biomass was extracted at a ratio of 1g of biomass to 20 mL of methanol for 24 hours at room temperature in stirred reaction vessels, and filtered through a Büchner funnel whereafter a second extraction took place for a further 24 hours. The filtrate from the second extraction was then combined with the filtrate from the first extraction and the methanol was removed using a rotary evaporator. The extract was then suspended in sterile water and spray dried using a nitrogen electrostatic spray dryer (FluidAir, Chicago, Illinois, USA) at conditions of inlet temperature -120˚C, outlet temperature -80˚C.

*PME Analysis:* A Waters Acquity H-Class UPLC and Xevo TQ-S Micro MS and Waters HSS T3 1.8 µm column (2.1 mm x 50 mm) (Waters Corporation, Milford, MA, USA) was used for determining tryptamine concentrations. A seven-point standard curve (10 ppb -1 ppm) was used to calculate concentration from the detector response. It was produced using analytical standards produced by Cerilliant (Round Rock, TX, USA) (psilocybin and psilocin) and Usona (Madison, WI, USA) (norpsilocin, baeocystin, norbaeocystin, and aeruginascin). Values obtained are given in the section, Drugs (above).

**Plasma psilocin levels**

To be certain that observed differences between the effects of PSIL and PME are not due to differences in plasma psilocin, we measured psilocin levels in the plasma of mice administered with these two treatments. Plasma psilocin levels after administration of PSIL 4.4. mg/kg i.p were (mean±SD) 308.1±29.0 (15 minutes), 308.9±31.5 (30 minutes) and 197.0±105.3 (60 minutes). After PME (PSIL dose 4.4 mg/kg i.p.) plasma psilocin levels were 344.4±79.1 (15 minutes), 283.7±62.2 (30 minutes) and 171.9±92.2 (60 minutes). No differences were statistically significant (Supplementary Fig. S38)

**Dose calculation for PME**

For PME the dose was based on the following calculation: To prepare a solution for a mouse that weighs 30 g so that the mouse will receive 4.4 mg/kg of pure psilocybin, $0.03\ kg\ (since\ 30\ g\ =\ 0.03\ kg)\ \times 4.4\ mg/kg = 0.132\ mg$ of psilocybin per 30 g mouse is needed. Since 1 mg of psilocybin is found in 76.92 mg of PME (1.3%) the amount of PSIL that was given needs to be multiplied by a factor of 76.92, $76.92 \times 0.132\ mg = 10.15\ mg$ of PME is needed per 30 g mouse. In all cases, injections were administered in a standard injection volume of 10 µl/g per mouse. Both PSIL and PME rapidly dissolved completely in saline (0.9% NaCl).
